# Numerical cognition based on precise counting with a single spiking neuron

**DOI:** 10.1101/662932

**Authors:** Hannes Rapp, Martin Paul Nawrot, Merav Stern

## Abstract

Insects are able to solve basic numerical cognition tasks. We show that estimation of numerosity can be realized and learned by a single spiking neuron with an appropriate synaptic plasticity rule. This model can be efficiently trained to detect arbitrary spatio-temporal spike patterns on a noisy and dynamic background with high precision and low variance. When put to test in a task that requires counting of visual concepts in a static image it required considerably less training epochs than a convolutional neural network to achieve equal performance. When mimicking a behavioral task in free flying bees that requires numerical cognition the model reaches a similar success rate in making correct decisions. We propose that using action potentials to represent basic numerical concepts with a single spiking neuron is beneficial for organisms with small brains and limited neuronal resources.

## 1. Introduction

Insects have been shown to possess cognitive abilities (Chittka and Niven, 2009; Avargùes-Weber et al., 2011, 2012; Avargùes-Weber and Giurfa, 2013; Pahl et al., 2013). These include estimating numerosity (Rose, 2018; Skorupski et al., 2018), counting (Chittka and Geiger, 1995; Dacke and Srinivasan, 2008; Menzel et al., 2010) and other basic arithmetical concepts (Howard et al., 2018, 2019). How insects succeed in these cognitive tasks remains elusive. A recent model study by Vasas and Chittka (2019) suggested that a minimal neural circuit with only four rate-based neurons can implement the basic cognitive ability of counting visually presented items. The study implies that their minimal circuits can recognize concepts such as a “higher” or “lower” item number and “zero” (Howard et al., 2018) or “same” and “different” number of items (Avargùes-Weber et al., 2012) when combined with a sequential inspection strategy that mimics the behavioural strategy of insects during detection (Dacke and Srinivasan, 2008). The neural circuit studied in Vasas and Chittka (2019) was shown to successfully predict whether a particular feature (e.g. yellow) has been presented more or less often than a pre-defined threshold number, despite being presented in a sequence of other features and distractors. This circuit model was hand-tuned in order to successfully estimate numerosity in a numerical ordering task similar to Howard et al. (2018). This poses the question how an efficient network connectivity could be learned by means of synaptic plasticity.

Numerosity estimation tasks that require to count the number of detected instances have also been researched in the field of computer vision, in particular in relation to object recognition tasks. Many resources have been devoted to train artificial neural networks to perform such tasks. Deep learning methods (Schmidhuber, 2015) in particular have been shown to be successful in object detection and they enable counting by detecting multiple relevant objects within a static scene either explicitly (Ren et al., 2015) or implicitly (Lempitsky and Zisserman, 2010). However, these model classes are costly as they typically need to be trained on a very large number of training samples (in the millions) and often require cloud-computing clusters (Krizhevsky et al., 2012; Simonyan and Zisserman, 2014). Indeed, OpenAI could recently show that the amount of computing power consumed by such artificial systems has been growing exponentially since 2012.

Clearly, insects with their limited neuronal resources cannot afford similar costly strategies but have to employ fundamentally different algorithms to achieve basic numerical cognition within a realistic number of learning trials. These biological algorithms might proof highly efficient and thus have the potential to inform the development of novel machine learning (ML) approaches.

A number of recent studies managed to train spiking neural networks with gradient-based learning methods. To overcome the discontinuity problem due to the discrete nature of action potentials, some studies evaluated the post-synaptic currents in the receiving neurons (Nicola and Clopath, 2017) and (Huh and Sejnowski, 2017) for the training procedures. Other studies used the timing of spikes as a continuous parameter (Bohte et al., 2000; O’Connor et al., 2017; Zenke and Ganguli, 2018), which lead to synaptic learning rules that rely on the exact time interval between spikes emitted by the presynaptic and spikes emitted by the postsynaptic neuron. These Spike Timing Dependent Plasticity (STDP) rules had first been observed experimentally (Bi and ming Poo, 2001) and since have gained much attention in experimental and theoretical neuroscience (Caporale and Dan, 2008; Song and Abbott, 2000). Other recent studies approached the problem by either approximating or relaxing the discontinuity problem (Zenke and Ganguli, 2018; Bengio et al., 2013) to enable learning with error backpropagation in spiking neural networks. Training single spiking neurons as classifiers has been proposed by Gütig and Sompolinsky (2006) and Memmesheimer et al. (2014). Closely related, Huerta et al. (2004) trained binary neurons to perform classification in olfactory systems.

Here, we study a biologically realistic spiking neuron model with a synaptic learning rule proposed by Gütig (2016). Our approach to numerical cognition takes advantage of the discrete nature of action potentials generated by a single spiking output neuron. The number of emitted spikes within a short time period represent a plausible biological mechanism for representing numbers. In a virtual experiment we train our neuron model to count the number of instances of digit 1 within a static image of multiple handwritten digits (LeCun and Cortes, 2010). The synaptic weights are learned from the observations and thus our model overcomes the problem of hand-tuning a single-purpose neuronal circuit. We then test the model on the same “greater than” task as in Vasas and Chittka (2019), but we use the model’s ability of precise counting to derive the concept of “greater than”.

Since in the present work we are interested in estimating numerosity, the teaching signal in our model represents a single integer value that equals the total number of relevant objects. To achieve successful training we introduced an improvement to the implementation in Gütig (2016). This approach presented in Gütig (2016) overcomes the spiking discontinuity problem by considering the membrane potential for the gradient-based learning. We show that our improved approach allows to train the model with better generalization capabilities and also supports better the reliability of numerosity estimation under inputs with complex distributions, including noise distributions, as naturally present in the brain.

## 2. Results

Our objective is the implementation of a spike-based method that can be trained to solve numerical cognition tasks. We employ the Multi-Spike Tempotron (MST) (Gütig, 2016), a single leaky integrate-and-fire neuron model with a gradient-based local learning rule. We suggest a modified update rule of the learning algorithm that reduces the variance in training and test error. The model is subjected to three different tasks that progress from a generic spike-pattern detection problem to a biologically inspired dual choice task that mimics behavioral experiments in honeybees.

### Detection of spatio-temporal input spike patterns

In a first approach we consider the problem of detecting different events over time. A particular event is represented by a specific spatio-temporal spike pattern across a population of neurons that are presynaptic to the MST. These spike patterns are generic representations of events that could, for instance, represent a sensory cue in an animal’s environment.

We generated event-specific patterns of fixed duration (1 s) across 500 presynaptic input neurons using a gamma-type renewal process of fixed intensity (*λ* = 0.89 spikes per second) independently for each neuron (see Methods). The MST was presented with an input consisting of a sequence of different patterns on top of a noisy background that was simulated as independent gamma-type renewal processes of either constant or time-varying intensity (see Methods).

A single input trial is shown in Figure 1A. It accounts for the random occurrence of three different event-specific spatio-temporal spike patterns (in this specific example, each pattern occurring once) as indicated by different spike color, and of distractor patterns occurring twice (black spikes). Gray spikes represent the background noise. Generally, for each trial of 10 *s* duration we randomly drew a number of pattern occurrences and pattern identities from a total of 10 possible patterns (5 target patterns and 4 distractor patterns).

**Figure 1:**
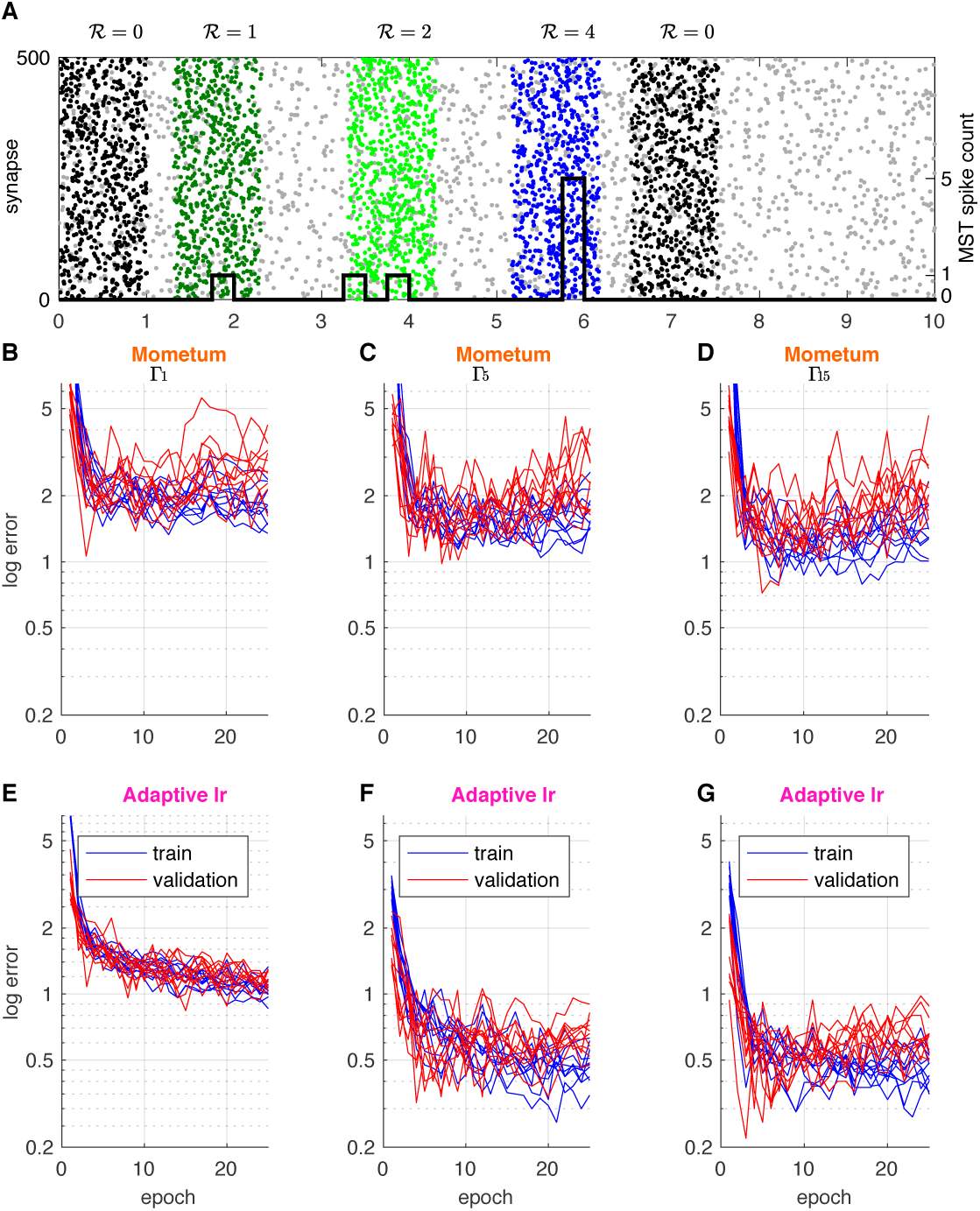
Comparison of training convergence for momentum and adaptive learning under different background noise conditions. **A.** Sample input sequence: A 10sec duration spike train input example. The spike train is composed of three patterns, each with a distinct target (dark green, green, blue), background activity (gray) and two distracting patterns (black). Number of MST output spikes superimposed as black step function. The MST is supposed to fire Σ*_i_*𝓡*_i_* = 7 spikes over the whole sequence, 𝓡 = 0 spikes for distractors and 𝓡 ∈ {1, 2, 4} for the colored patterns. Patterns are simulated with gamma processes of different order (separate data-sets): Γ_1_ (Poisson), Γ_5_, and Γ_15_. Patterns are superimposed onto 10sec inhomogeneous Poisson background activity. **E-G.** Learning curves (blue) and validation curves (red) for 10 independent simulations of the **B,E.** Γ_1_ (Poisson), **C,F.** Γ_5_, and **D,G.** Γ_15_ patterns. **B-D.** MST with momentum-based learning implementation (Gütig, 2016). **E-G.** MST with adaptive learning implementation. Learning convergence shows larger variance when using momentum as compared to using adaptive learning. The same is true for the validation error. This indicates that adaptive learning is capable of finding better optima as compared to momentum.

We first trained the original MST of Gütig (2016) to detect pattern occurrence. To each of the 5 event-specific patterns we assigned a specific target number of MST output spikes 𝓡 (from 1 to 5) and the MST should produce zero output spikes for any distractor pattern. At the end of each training trial the sum of actual output spikes is evaluated and compared to the desired number of output spikes determined by the trial-specific random realization of the input pattern sequence. The absolute difference between desired and actual spike count determines the training error in the range of 0 − *N* ∈ ℕ_+_. If the actual number of spikes is larger than the sum of desired target spikes by some Δ*_k_*, a training step of the MST is performed towards decreasing its output spikes by the difference Δ*_k_*. Similarly, if the actual number is smaller than the sum of desired target spikes, a training step is performed to increase the MST’s number of output spikes by Δ*_k_*. No training step is performed for correctly classified samples.

To analyze model performance we computed the training error and validation error for up to 25 training epochs (see Figure 1B-D). Each training epoch consisted of a fix, randomized set of 200 trials, the validation set consisted of 50 trials. Both, training error (blue) and validation error (red) dropped sharply with increasing number of training epochs and reached a plateau at about 2 spikes after ∼ 10 epochs, independent of the type of the gamma-order used for pattern generation (Figure 1 B-D).

### Local synaptic update method improves performance and robustness

Training and test errors exhibited a high variance across repeated models (Figure 1 B-D) indicating limited robustness of model performance. We therefore replaced the Momentum method for gradient descend implemented in the original work of Gütig (2016) by a synaptic specific adaptive update approach similar to RMSprop as proposed by Tieleman and Hinton (2012) (see Methods).

While speed of convergence is similar when using the adaptive learning method compared to Momentum, we find that using adaptive learning results in less variant training error (Fig. 1E-G and Fig. 2A,B). This also holds for the variance of the test error on an independent validation set indicating better generalization capabilities to previously unseen inputs. The adaptive, per synapse learning rate combined with exponential smoothing over past gradients has a regularizing effect and prevents the model from over-fitting to the training data. We further conclude, that the modified algorithm is potentially able to find better and wider optima of the error surface as compared to learning with Momentum. More importantly this behaviour is consistent and independent of the spike generating process and noise level (fig. 2A,B).

**Figure 2:**
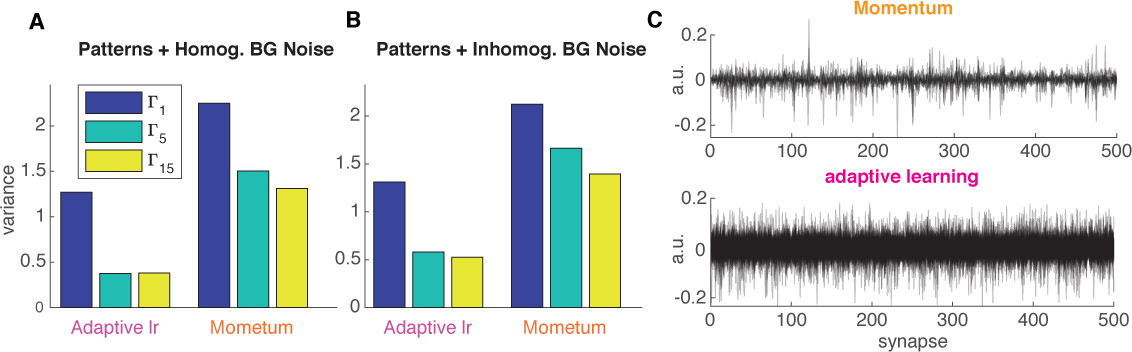
**A,B** Variance of validation error measured at epoch 10 for data sets with homogeneous (**A**) and inhomogeneous (**B**) background noise. **C:** Empirical analysis of the regularizing effect on the error variance. Weight changes Δ*ω_i_* over all training steps (and all epochs) are collected for each synapse *ω_i_*. PCA is performed to reveal, which synapses show the largest variance of Δ*ω_i_* over the entire training process. Large variance in Δ*ω_i_* implies strong modification of a synapse. For both methods the first 10 principal components are shown where *x* axes correspond to the synapses *ω_i_* and *y* axis can be interpreted as the magnitude of total weight change during the training process applied to each synapse. The Momentum method tends to tune only a small subset of the available synapses strongly whereas the adaptive learning method leads to modifications that are more uniformly distributed over all synapses and more broadly distributed in magnitude.

At this point we cannot provide a theoretically grounded explanation for the regularizing effect we see when using adaptive learning instead of Momentum. Development of theoretically grounded explanations of the effects of different gradient-decent optimizers is a very recent and active research field in the Deep Learning community. Even for rate-based artificial neural networks it’s currently not possible to provide a sound theoretical explanation. We conducted an empirical analysis of the weight updates shown in Figure 2C. Specifically, we performed PCA on the weight changes Δ*ω_i_* applied to all synapses over all training steps. The intuition here is that large variance in Δ*ω_i_* implies strong modification of a synapse over the training process. Results of our analysis (Figure 2C) show that for the adaptive learning method the weight changes are more uniformly distributed over all synapses and more broadly distributed in magnitude. In contrast, with the Momentum method only a small subset of synapses are strongly modified. We conclude that distributing the updates uniformly over all synapses leads to a more deterministic convergence behavior towards good minima in the error surface, independently from the initial, random initialization of *ω_i_*. The results shown are obtained from a specific choice of meta-parameters (*α* = *γ* = 0.99, *λ* = 0.01) but we verified that it remains true over a broad range of possible values & combinations.

Moreover, we find that adaptive learning improves absolute performance converging to a smaller error independent of the actual gamma process when using the same values for the free meta-parameters for both methods. While choosing different values for the meta-parameters result in different (and in some cases even lower) train and validation errors, our main result regarding the variance still holds. For subsequent tasks we used the MST with adaptive learning.

### Counting handwritten digits

We apply the MST model to the problem of counting the number of instances of digit 1 within an image showing several random handwritten MNIST digits (LeCun and Cortes, 2010). The digits are randomly positioned within a fixed 3 × 3 grid (Figure 3A). Each image can contain between 0 and 6 instances of the digit 1 at one of the 9 possible grid locations. To solve this problem with the MST we take the 50*x*50px input image and encode the entire image as parallel spike train. To transform the image into a parallel spike train that can be fed into the MST model we use Filter-overlap correction (FoCal) of Bhattacharya and Furber (2010). This method is an improved 4-layer model of the early visual system using rank-order coding as originally proposed by Thorpe and Gautrais (1998). The MST model is then trained to count the number of occurrences of digit 1 by generating one output spike for each instance of digit 1 (Figure 3A). The MST is trained on targets 0 − 5 using 5-fold cross-validation on 400 sample images. The learning rate is tuned manually to *λ* = 0.00002 which yields the best performance and training speed. For reference we compare the performance of the MST to a conventional computer vision model that uses a convolutional neural network (ConvNet) (Krizhevsky et al., 2012; Seguí et al., 2015). The ConvNet is trained similarly but provided 800 training samples and a larger learning rate of 0.01 to speed up the training process.

**Figure 3:**
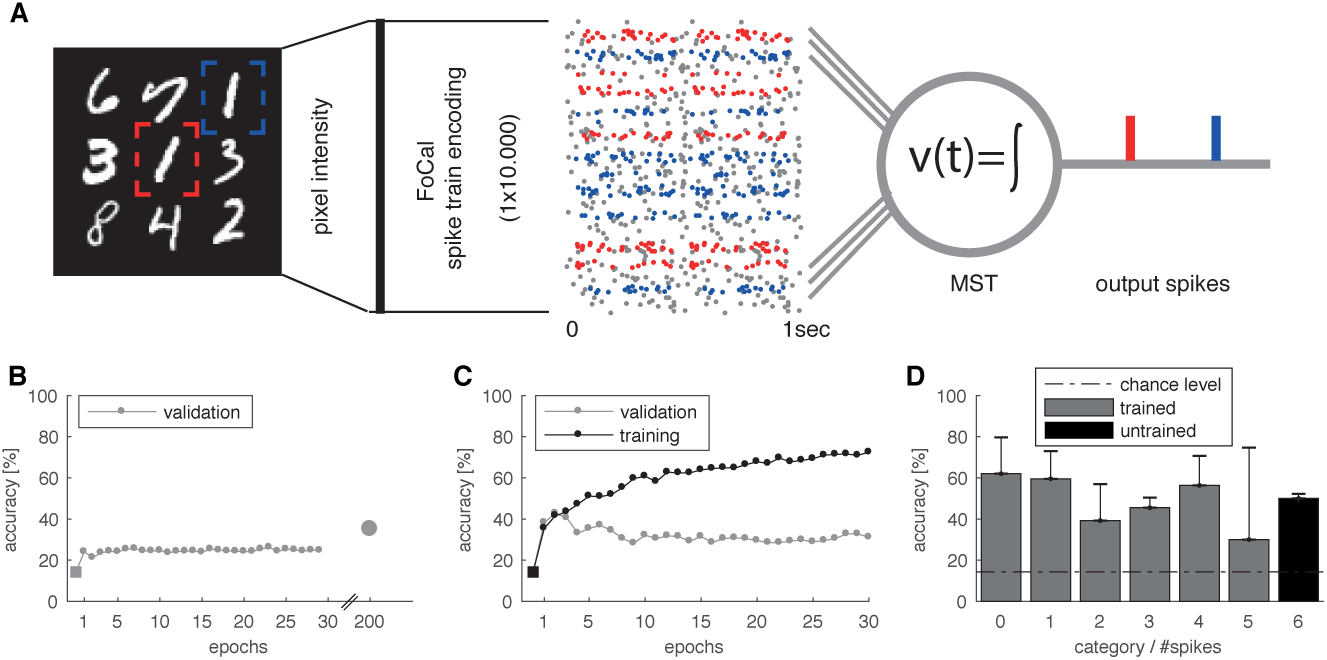
Counting of visual concepts with spikes. **A:** Sketch of counting task. Goal of this task is the accurate estimate of the number of occurrences of digit 1 in an image of random MNIST digits. Example image (50 × 50 px) with multiple random digits from the counting MNIST data set positioned within a 3 × 3 grid. The image is encoded into parallel spike trains by applying FoCal encoding, resembling a 4-layer early visual system with rank-order coding. The multivariate spike train converges onto the MST via 10.000 plastic synapses. The MST is trained to elicit exactly *k* output spikes where *k* is equal to the number of digit 1 occurrences in the original image (here 2). **B:** For reference we trained a ConvNet on the same raw images. Shown is the performance in terms of mean accuracy (5-fold cross-validation). After 200 training epochs the ConvNet reached ~ 40% accuracy. **C:** Performance of the MST in terms of mean accuracy (5-fold cross-validation). The MST shows rapid learning reaching a similar level of accuracy as the ConvNet after 200 training epochs within only 2-4 training epochs. **D:** Mean accuracy + std for the possible numbers of digit 1 present within a single image (categories). The MST is trained on samples of categories 0 − 5 to generate 0 − 5 output spikes respectively. The MST is then tested on the untrained category 6 and is able to generalize reasonably while the ConvNet, by design, cannot make predictions for this category.

Counting, as a conceptual problem, is similar to a regression problem where we have no a-priori knowledge of the maximum number of desired targets present in an input. It is important to note that the ConvNet model used for comparison is built using prior knowledge about the distribution of the training set. The ConvNet is constrained to learn a categorical distribution over [0, 5], where 5 is the maximum possible count of desired digits in the used training set of images. This has two implications. First, the ConvNet model will be unable to predict images that include more than 5 targets. In general for regression problems the prediction targets are usually not bounded. Second, the counting error a ConvNet can make is constrained by the training bound, i.e. the maximum error is 5. In contrast, the MST model does not have any need for this prior knowledge or constraints. In principle it is capable of solving the general, true regression problem and can (after being trained) also make predictions for images that contain more than 5 occurrences of digit 1. It thus has to solve a more difficult learning problem. The maximum prediction error in this case is unbounded rendering the MST more vulnerable to prediction errors compared to the ConvNet. Figure 3B shows the performance of the ConvNet in terms of mean accuracy of correctly counted images. Despite the large learning rate, accuracy only slowly (but monotonically) improves over the course of 200 training epochs. In contrast, the performance of the MST in Figure 3C shows rapid learning, reaching similar mean accuracy as the ConvNet within only ∼ 3 training epochs. The MST reaches a performance above chance level for each of the trained target categories 0 − 5 (Fig. 3 D). It also performs above chance level for images that contain 6 targets. This indicates that the MST is not only learning a categorical distribution over 0 − 5, as is the case for the ConvNet, but generalizes to a larger, previously unseen number of targets. We want to emphasize that the MST performs better than the ConvNet despite the advantages given to the latter in form of a larger number of training samples (200% of the samples given to the MST) and a higher learning rate (by factor 1000). The results are further summarized in table 1.

**Table 1:** Results for MNIST digit counting MNIST task. We evaluate each model in terms of root mean square error (RMSE) of the difference in actual and predicted number of digits (lower is better) and accuracy of correct digit count in images. Reported results are mean and standard deviation over a 5-fold cross-validation.

| Counting MNIST results: counting ones |  |  |  |
| --- | --- | --- | --- |
| Model | #Parameters | RMSE (mean ± std) | Accuracy (mean ± std) |
| ConvNet (5 epochs) | 11079 | $1.70 \pm 0.2083$ | $23.00 \pm 0.1746$ |
| ConvNet (100 epochs) | 11079 | $1.67 \pm 0.3130$ | $27.07 \pm 0.5113$ |
| ConvNet (200 epochs) | 11079 | $1.02 \pm 0.0768$ | $40.97 \pm 0.8136$ |
| **MST<sub>adaptive</sub> ($\sim 4$ epochs)** | 10000 | $1.21 \pm 0.2067$ | **$47.72 \pm 3.2052$** |

During our experiments we found that the choice of the spike encoding method has a big impact on the MST’s performance. It is possible that, by applying better or more efficient encoding algorithms, the performance of the MST model can be further improved.

### Insect-inspired numerical cognition during visual inspection flights

We now consider a biologically motivated task following Vasas and Chittka (2019) and the original experiment conducted in honeybees by Howard et al. (2018). The objective in this experiment is to perform a ‘greater than’ dual choice task on two stimulus images that show varying numbers of geometric shapes (circles, squares, diamonds). The geometric shapes within a stimulus image are consistent and the possible number of geometric items present are 1 − 6. In contrast to our previous task, here a stimulus image is not presented as single static input. Instead the input is a sequence of smaller images that relate to a 60° field-of-view (FOV) of honeybees hovering over the stimulus image at a distance of 2 cm (see Methods). The available data set is highly imbalanced and limited to a total of 97 stimulus images and corresponding inspection flight trajectories recorded from behaving honeybees. Figure 4A shows an example stimulus image with 6 diamond shapes and the inspection trajectory taken by one honeybee. This particular trajectory yields a sequence of ∼ 40 FOV images (red dots). Following the exact same procedure as Vasas and Chittka (2019), the (absolute) derivative |*S*(*t*) − *S*(*t* + 1)| of two subsequent FOV images *S*(*t*), *S*(*t* + 1) is computed as input to the model (see Figure 4B). To reduce computational cost for our MST model and to unify the varying sequence length across all stimuli we sub-sample the trajectories to length 10 (magenta dots). In Vasas and Chittka (2019) a rate-based model was used and the FOV images were encoded into a univariate time-series (representing a rate) that is fed into the model as a single presynaptic input. Since our MST is a spiking model we have to encode each FOV image into a spike train. We apply the same encoding strategy as used before in the counting MNIST task: Each FOV image is encoded as a parallel spike train of 10000 synapses using FoCal. All encoded FOV images are combined into a long parallel spike train by concatenation (see Figure 4C).

**Figure 4:**
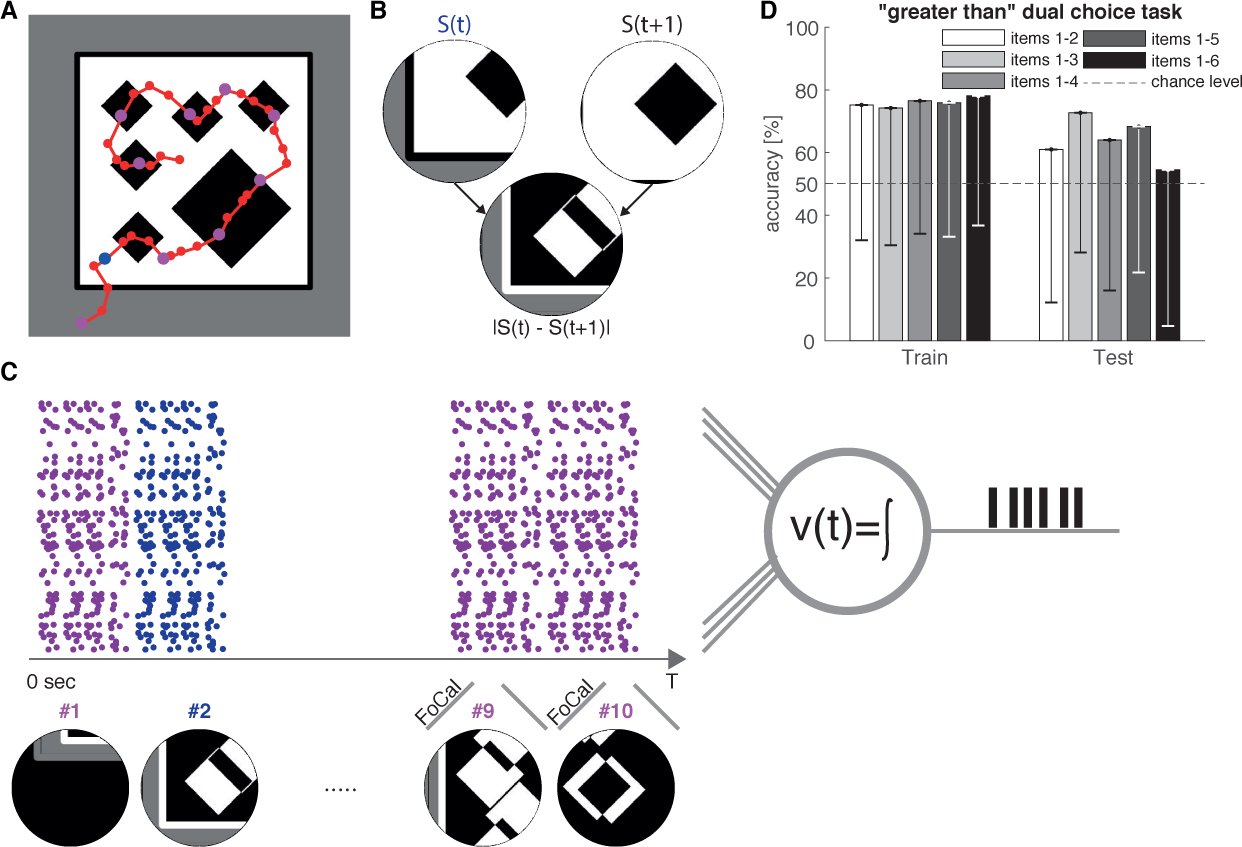
Dual choice ‘greater than’ task performed on geometric shapes using a visual inspection strategy observed in honeybees. **A:** Sample stimulus image with 6 diamond shapes and inspection trajectory (red) of a honeybee. **B:** Visual input *S*(*t*) perceived by the honeybee during the inspection trajectory (red curve, sampled at 40 time points) with a circular field-of­ view (FOV) of 60°. Following the method of Vasas and Chittka (2019), input to the model is constructed as derivative of two subsequent FOV images from two subsequent time points: *FOV_diff_* = |*S*(*t*) − *S*(*t* + 1)|. **C:** Sequences of FOV images are constructed by sub-sampling the inspection trajectory to 10 FOV images. For each of the 10 time points the *FOV_diff_* is encoded into spatia-temporal spike patterns using rank-order coding (FoCal) and concatenated (without gaps) into the resulting parallel spike train. The MST is trained (supervised) to match its output spikes to the number of geometric items in the original stimulus image shown in panel A (precise counting). **D:** Performance in the ‘greater than’ dual choice task. The MST output (number of spikes) in response to 2 different stimulus images is used and compared. When 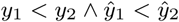 the decision is considered correct (and vice versa for *y*_1_ > *y*_2_)· Bars show mean accuracy - std and grouped by increasing maximum number of items present per image. Our results indicate that the MST can achieve mean accuracy that is comparable to that of honeybees reported in Howard et al. (2018)

The task is divided into two steps. The MST counts the number of geometric items present in a stimulus image. The resulting count numbers are then compared to solve the ‘greater than’ dual choice task. This differs from the original behavioral task Howard et al. (2018) in which the honeybees were directly trained on the ‘greater than’ decision rather than on precise counting.

To this end we trained the MST (using 10-fold cross-validation) to match the number of generated output spikes to the number of geometric items present. To evaluate the dual choice task we take 2 random stimulus images with a different number of items *y*_1_, *y*_2_ and feed these images as input into the trained MST. We then compare the true item count with the predicted item count 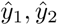 of the MST. If 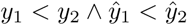 it’s considered a correct decision and vice versa when *y*_1_ > *y*_2_. For undecidable cases where 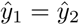 a random decision is taken. This sampling process of decisions is repeated for 1000 iterations. Our results (Figure 4 D) show that the MST model is able to achieve comparable performance to the average performance of the honeybees (60 − 70%) in the original task of Howard et al. (2018) in terms of mean accuracy of correct decisions. We want to emphasize that the MST performance could be achieved despite the very small and imbalanced training data. Moreover, the MST is trained on the problem of precise counting that is harder than the binary decision task. While we have to acknowledge that the results show large error bars (due to the very limited training data) we conclude that our results provide a successful proof-of-concept. Using a larger and more balanced training set and better feature encoding would certainly reduce the variability and further improve the performance.

## 3. Discussion

### Counting as a basis for numerical cognition

Numerical cognition is a general term that covers several sub-problems, for example numerosity, counting, relations (greater/smaller than), basic arithmetical operations and many more. While each individual sub-problem might appear fairly trivial to us as humans, it’s not clear yet how this could be realized computationally on the level of spiking neurons or networks thereof. Despite the simplicity of these sub-problems they do provide a foundation for more complex concepts that humans make heavy use of and are relevant for behavioral decision making. For example, if one is able to count entities it might only be a small step towards combining that information to perform more advanced concepts such as empirical statistics and estimating (discrete) probabilities. While the specific symbolic math concepts are unavailable to animals, they are still able to show basic numerical cognition and to evaluate basic probabilities (Howard et al., 2018; Avargùes-Weber et al., 2012; Howard et al., 2019).

A first objective of the present work was to study whether a single neuron model has the computational power to support numerical cognition tasks. Specifically, we addressed cue detection and counting by adjusting neuronal input such that it generates an output spike count that matches the number of relevant cues in its input. In order to achieve this computationally, the presynaptic weights of the neuron need to be tuned. Given the fact that the parameter space is very large and many possible solutions may exist, manually tuning the parameters is usually not possible. It is therefore desirable to implement a plasticity mechanisms that allows the neuron to tune its weights by learning from examples.

In this work we have explored the Multi-Spike Tempotron (MST) developed by Gütig (2016). This spiking neuron model can be trained by gradient-descent to produce a precise number of output spikes in response to multiple occurrences of patterns embedded in the presynaptic input. Different patterns are assigned to different target numbers of output spikes per pattern occurrence. Gütig (2016) showed that the MST is capable to learn to detect different spike input patterns. It can further assign the correct number of output spikes matching the targets of individual patterns. The MST learns this despite the fact, that the (supervised) teaching signal is only provided as a single scalar value that is equal to the sum over all targets presented sequentially in the input. Departing from the homogeneous Poisson process studied in Gütig (2016) we confirmed MST performance for biologically more realistic gamma processes as generators for input patterns on non-homogeneous background.

### Adaptive local learning rule benefits model robustness

We introduced a modification to the update rule of synaptic weights Δ*ω_i_*. The *adaptive learning* introduces a dynamic, synapse-specific learning rate whose value at training step *t* depends on its history of values from previous training steps. We find that this modification allows the MST to learn a parameter set for the synaptic weights that shows less variability of the training and validation error as compared to the original *Momentum* method used in Gütig (2016). Low variability in validation error is generally a desired property for any learning algorithm since this commonly implies low variability in prediction or classification performance on new, unseen data.

At this point we are unable to provide a theoretically grounded explanation of the regularizing effect shown by *adaptive learning*. The deep learning community currently still lacks theoretical understanding of the effects of different gradient-descent optimizers, which is actively researched (Choromanska et al., 2015; Jin et al., 2017). We performed an empirical analysis of the weight changes Δ*ω_i_* over the course of training. Specifically, we used PCA to analyze the variance of Δ*ω_i_* for each synapse over all training steps. Our analysis reveals that for the *adaptive learning* a large number of weights is affected. In contrast, when using the originally proposed *Momentum* method, a much smaller subset of synapses show significant weight changes and their distribution appears much more heavy-tailed with strong weight changes in few neurons. We conclude that modifying all synapses uniformly appears to increase the likelihood that training converges more deterministically towards good minima, independent from the initial random choice of *ω_i_*.

### Spike-based biological learning versus machine learning

A second objective of this work was to explore possible advantages and disadvantages of a spike-based learning algorithm in comparison to a state-of-the-art deep learning architecture. Biological learning mechanisms enable animals to learn rapidly in a complex and dynamic environment. Instances where sensory cues coincide with reward or punishment during exploration may be sparse, i.e. they have to learn on very small sample sizes and slow learning could have fatal consequences. Single-trial learning, for instance, seems to be a fundamental ability found in lower and higher animals. Insects, for example, are able to form long-lasting associative memories upon a single coincident presentation of a sensory stimulus and a reinforcing stimulus (Scheunemann et al., 2013; Pamir et al., 2014; Zhao et al., 2019). Increased learning speed generally comes at the cost of increased generalization error and thus learning speed and high accuracy are in trade off.

We therefore compared the biologically inspired spike-based learning algorithm of the MST to the deep learning architecture of a convolutional neural network, a standard computer vision model (ConvNet). We found that the MST is able to rapidly learn this task within ∼ 3 training epochs of 320 samples each to achieve a mean accuracy of ∼ 47% of correctly counted digits. Additional training did not improve accuracy. Conversely, the ConvNet, despite a 1000× larger learning rate and 100% more training samples per epoch, required > 200 training epochs to achieve a similar accuracy. With additional training, the ConvNet achieved > 80% accuracy for > 5000 training epochs (not shown). Our results reflect a trade-off between very fast but less accurate learning with the spike-based MST method versus slow but increasingly accurate learning with the ConvNet. An additional aspect of biological relevance is the consumption of (computing) power that is considerably higher for the ConvNet than for the single neuron MST. From the biological perspective, processing with spikes is generally energy efficient, an important constraint in living organisms (Levy and Baxter, 1996; Niven and Laughlin, 2008; Niven, 2016).

Once trained, the ConvNet is only able to learn a categorical distribution over a fixed set of possible targets (here 0 − 5) that needs to be put into the design of the model a-priori. Similarly, previous related work of gradient-based learning in spiking network models are mostly concerned with solving classical classification tasks with pre-defined classes (Zenke and Ganguli, 2018; Bohte et al., 2000; Gütig and Sompolinsky, 2006; Memmesheimer et al., 2014). In this work we applied the single-neuron MST model to solve a regression problem. We show that the MST model does not have the limitation of the ConvNet. After being trained on targets 0 − 5 it was able to generalize to previously unseen images that contained digits 1 at 6 out of the 9 possible positions. This indicates that, in principle, the MST can solve full regression problems.

Differently from all other tasks presented in this work, the difficulty in this task is that each input stimulus is presented as a whole and not sequentially. This means that spike patterns associated with each occurring instance of digit 1 are distributed spatially (over different sets of synapses) instead of temporally. Due to the random positioning within the 3 × 3 grid the patterns to be identified by the MST ‘*jump*’ over different sets of synapses for different stimulus images that share the same training target, which makes this task particularly hard to solve.

### 3.1. Relational operation based on counting

In the final task we went one step further and studied the hypothesis that (precise) counting allows other basic numerical cognition tasks to emerge. Assuming that a single neuron can count by relating the sum of its output spikes to the number of items present in a single stimulus, we show that this allows to solve other numerical cognition tasks. To this end we use a biologically motivated ‘greater than’ dual choice task, performed by honeybees that employ a sequential inspection strategy. Honeybees are presented with stimulus images that show 1 − 6 different geometric shapes. Given 2 different stimulus images, the bees have to decide which of the two images contain more geometric items. Due to their limited field-of-view (FOV), the bees cannot perceive the stimulus image as a whole. Instead they perform a sequential inspection strategy, by hovering over the entire stimulus image. This results in a time series of FOV images, similar to a moving spot light. Using the MST, we approach this problem similarly and present a long parallel spike train that contains a sequence of FOV images. In contrast to the honeybees the MST is trained to perform precise counting of the geometric shapes, similarly to the counting task we presented earlier. To perform the ‘greater than’ dual choice task we present two different stimulus images and compare the number of output spikes of the MST. We show that the MST is able to achieve average success rates in terms of correct decisions that are comparable to those achieved by honeybees in the original experiment. While our results do show much larger error bars as the honeybees, this is due to the following important differences that need to be considered: The bees are explicitly trained on the (binary) ‘greater than’ task. Contrary, the underlying problem that the MST solves here is *precise counting*, which is harder to solve in general. Differently from Vasas and Chittka (2019), where input is provided as a univariate rate signal, our model uses parallel spike trains as input, that are derived from the FOV images. While, Vasas and Chittka (2019) used hand-crafted features based on the assumption that the number of step-like changes in global image contrast is proportional to the number of scanned items, the MST has to learn which relevant features to extract from the spatio-temporal input spike patterns during the training process. The MST has been trained on this task with a very small data set of ∼ 70 samples per trained model. Increasing the training data will very likely result in better and especially more robust performance and hence smaller error bars.

### Limitations of the Multi-Spike Tempotron

The MST learning rule used in this work requires differentiation of the membrane potential (see Methods), which is considered to be biologically implausible. Gütig (2016) suggested an approximate formulation of the learning rule that uses correlation-based learning of presynaptic spikes and postsynaptic voltage, which is considered biologically plausible. In order to ensure comparability of our results with the results in the original work we here used the gradient-based learning rule that was evaluated in Gütig (2016) for all the experiments presented. While in this work we specifically focused on the computational capabilities of the single-neuron model, the same model and learning rule could also be used to construct more complex and layered networks as shown in Gütig (2016). We leave the study of multiple interconnected MST neurons for future research.

One weakness of the MST identified in the course of our study is a tendency to overfitting. This can for instance be inferred from the insect-inspired numerical cognition task, where the MST can be trained to reach > 80% in accuracy of precise counting on the training set, but performance on the test set remains low or even drops below chance level (data not shown). This indicates that the MST tends to learn to memorize the samples of the training set instead of learning features that would generalize to the test set. This, to some extend, is also the case for the MNIST task. A potential solution to this problem could be to introduce explicit regularization terms in the MST learning rule, similarly to approaches realized in deep learning algorithms. During our experiments we further found that the choice of method for the multivariate spike encoding of images has a big impact on learning and prediction performance. We predict that improved or more sophisticated spike-encoding methods will boost performance.

### Conclusion

Action potentials represent an elemental discrete quantity of information processing in nervous systems. We conclude from our study that action potentials produced by a single spiking neuron can support the basic arithmetic operation of counting. The MST is a powerful single-neuron method that can be trained to solve regression problems on multivariate synaptic input. We successfully applied the MST to perform basic numerical recognition tasks on complex and noisy input. We suggest that using spikes to represent numerosity with a single neuron can be a beneficial strategy especially for small-brained organisms, which economize on their number of neurons.

## Acknowledgments

This work has been sparked during the OIST Computational Neuroscience Course and has been supported by accommodations and travel grants from the Okinawa Institute of Science and Technology (OIST) to HR. HR is supported by the German Research Foundation (grant no. 403329959 to MN) within the Research Unit ‘Structure, Plasticity and Behavioral Function of the Drosophila mushroom body’ (DFG-FOR 2705, www.uni-goettingen.de/en/601524.html).

## Author Contributions

Conceptualization, H.R., MP.N. and M.S.; Methodology, H.R and M.S.; Writing – Original Draft, H.R. and M.S., Writing – Review and Editing, H.R. and MP.N., Writing – Review and Editing of revised manuscript, H.R. and MP.N.

## Supplemental Information

### Transparent Methods

To support further research we make our code and data-sets publicly available at Rapp and Stern (2018).

### Multispike Tempotron Model

The Multi-Spike Tempotron (MST) is a current-based leaky integrate-and-fire neuron model (Gütig, 2016). Its membrane potential, *V*(*t*), follows the dynamical equation:

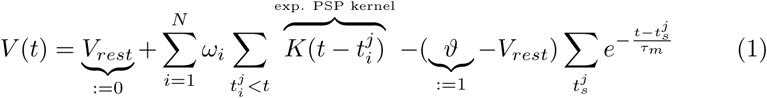

where 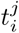 denotes the time of spike number *j* from the input source (presynaptic) number *i*, and 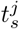 denotes the time of postsynaptic spike number *j* of the Tempotron neuron model. For mathematical convenience the resting potential is chosen to be *V_rest_* = 0 and *ϑ* = 1. Thus equation 1 can be simplified to:

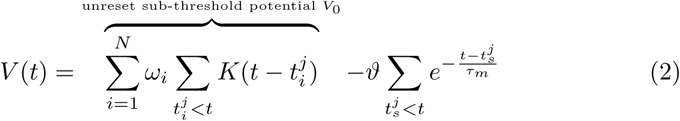

Every input spike at 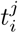 contributes to the postsynaptic potential (PSP) by the following causal kernel:

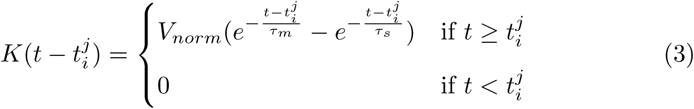

multiplied with the synaptic weight *ω_i_* of input synapse *i*. These synaptic input weights are learned via the gradient decent algorithm. The kernel is normalized to have its peak value at 1 with 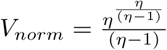 and 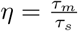 where *τ_m_* and *τ_s_* are the membrane time constant and the synaptic decay time constant. The kernel is made causal by setting it to 0 for 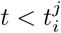. When *V*(*t*) crosses the spiking threshold *ϑ* the neuron emits a spike and is reset to *V_rest_* = 0 by the last term in equation 2 (soft-reset).

In order to have the neuron emit the required number of *k* postsynaptic spikes in response to some presynaptic spike pattern the weights *ω_i_* are modified. Since the required number of postsynaptic spikes are non-differentiable discrete numbers the gradient for adjusting the weights is derived from the spiking threshold using a continuous objective function, the spike-threshold surface (STS). The STS is a step function ℝ^+^ ↦ ℕ_0_, which maps each threshold value *ϑ* to the number of output spikes (*ϑ* ↦ *STS*(*ϑ*)) that will be generated by the neuron with this threshold value. The STS for a presynaptic input can be described by the decreasing sequence of critical thresholds values 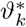:

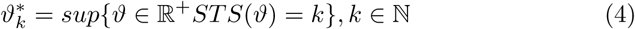

The critical threshold 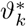 denotes the threshold value at which the neuron’s number of generated output spikes jumps from *k* − 1 to *k*. The number of generated output spikes remains constant when *ϑ* is between two critical threshold values: 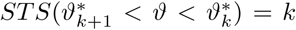. Additionally, a neuron does not fire any output spike if its threshold is larger than the maximum postsynaptic voltage (*V_max_*). In this case the STS is zero: *STS* (*ϑ* > *V_max_*) = 0. The first output spike is generated when *ϑ* = *V_max_*, thus the critical threshold for *k* = 1 spike is 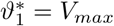. Generally, all 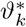 are just regular voltage values and can be described by the neuron’s membrane equation 2 which is a function of the synaptic weights *ω_i_* of the neuron. Hence, all critical thresholds are also a function of *ω_i_* and thus differentiable with respect to them. The goal is to adjust 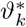 (by modifying the synaptic weights *ω_i_*) whenever the number of generated spikes does not match the desired training target. In our case, the specific *k* of desired output spikes is provided as supervised teaching signal. For each presynaptic input where the number of output spikes did not match the desired training target a training step is performed to adjust the number of output spikes towards *k*: Δ*k* = |*k_generated_* − *k_target_*| and *η* = *sign*(*k_generated_* − *k_target_*) indicates whether the neuron should increase or decrease its number of output spikes by Δ*k*.

To simplify notation, from now on we denote *ϑ\** as the desired critical threshold, e.g. 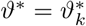 for the desired *k* of a specific presynaptic input.

The gradient of the critical threshold can be found by:

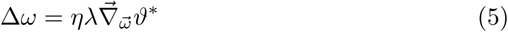

Where *η* ∈ {−1, 1} controls whether to increase or decrease the number of output spikes towards the *k* required spikes, *λ* is the learning rate parameter that controls the size of the gradient step to take in each training step and 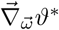 is the gradient of the critical spiking threshold with respect to the synaptic weights. To evaluate the expression in eq. 5 the properties of the critical spike time *t\** is used where by definition of the neuron equation 2 and *ϑ\** the following identity is satisfied:

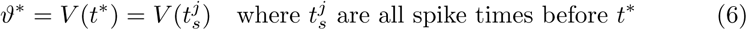

In what follows a recursive expression is derived for the gradient in equation 5 using equations 2 and 3. For notional clarity the recursive expression for the gradient is derived for a single component *ω_i_* of the vector 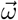. The generalization to 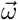 is immediate.

Let *m* denote the the number of output spikes the neuron fires before *t\**: 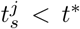 for *j* ∈ {1, …, m}. Using the identities in 6, for each synapse *i* the derivative of *ϑ\** has the following properties:

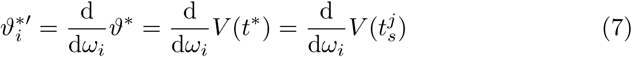

And the derivative of *ϑ\** follows the equation:

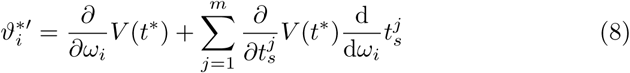

In the last equation the vanishing term 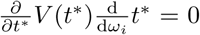 has been dropped. This relationship is true because *V*(*t\**) is either a local maximum with 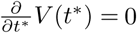 or *t\** is the arrival time of an inhibitory input spike that does not depend on *ω_i_*.

Similarly for each *k* ∈ {1, .., *m*} the following relationship holds:

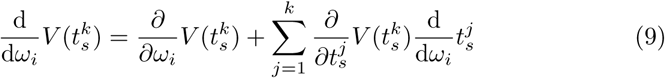

from which the following equations are obtained:

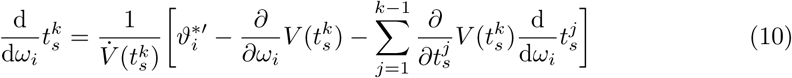

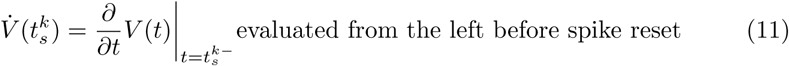

To solve equation 8 for 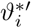, the right hand side of eq 10 is refactored to:

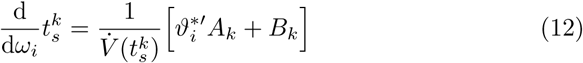

The coefficients *A_k_*, *B_k_* are given by the following recursive equations:

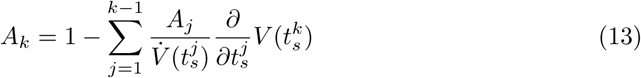

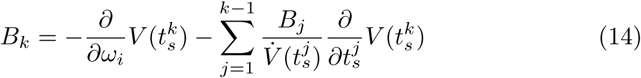

Similarly for *t*\* the analogous recursion formula is defined:

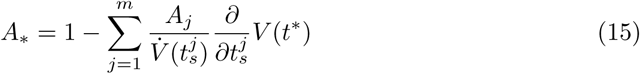

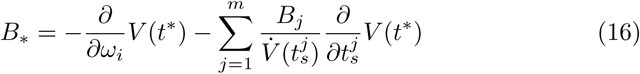

Inserting equation 12 into 8 the derivative of 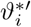 for each vector component *i* of *ω* can be expressed as:

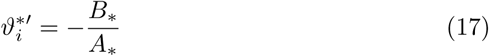

To calculate *A*_*_ and *B*_*_ all times 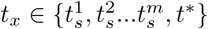 are considered at which the voltage reaches the spiking threshold *ϑ*. At these time points, due to the spiking and soft-reset, the membrane potential equation 2 reduces to the form:

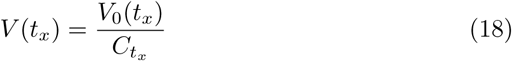

with

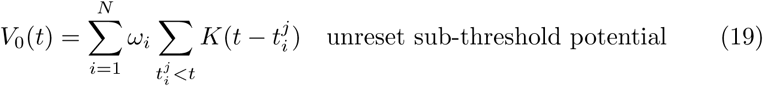

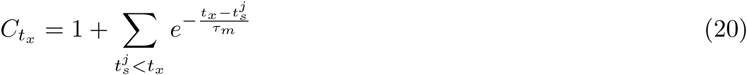

and gives the following derivatives:

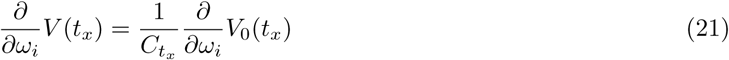

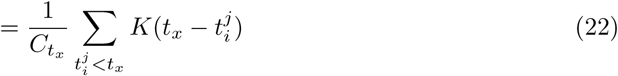

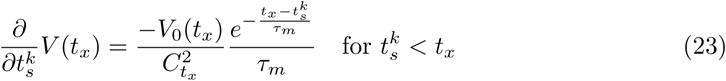

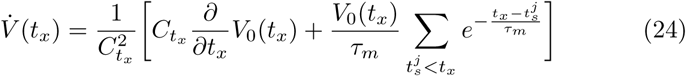

Where in our implementation the temporal derivative 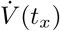 is estimated numerically instead of using its analytical expression.

### Momentum and Adaptive learning

The learning rate *λ* is global for all synaptic weights. Hence, the gradient descent takes an equal size step along all directions. If this parameter is too small the training process will take very long, but if it’s too big the algorithm might miss an optimum within the error surface and never converge to a good solution. Hence, tuning this learning rate is important to achieve decent training speed. A possible approach (Gütig, 2016) to avoid these problems is to update the weights according to exponential moving average of current and past gradients (up to training step t), using the *Momentum* heuristic:

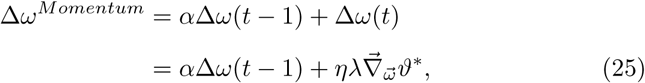

where *α* is the *Momentum* meta-parameter to control the exponential smoothing effect. In practice a common heuristic for the choice of *α* in the deep learning community is 0.999 and only tuning the global learning rate *λ*.

### Adaptive input weight learning and gradient smoothing

We propose here to use an adaptive learning approach for the weight updates instead of the *Momentum* heuristic. The proposed algorithm fits each input synapse with its own update rate and by doing so it takes into account that each synapse contributes to the overall update with a different level of importance. For example, updates should be larger for directions that provide more consistent information across examples. The RMSprop (Root Mean Square (back-)propagation) (Tieleman and Hinton, 2012) is a possible approach to achieve this. It was successfully used in deep learning for training mini-batches. It computes an adaptive learning rate per synapse weight *ω_i_* as a function of its previous gradient steps :

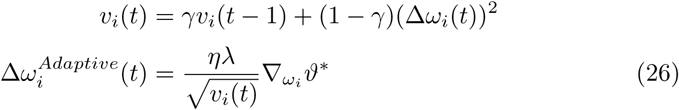

The dynamical variable *v_i_*(*t*) gives the synapse specific (e.g. local) learning rate for the current training step *t*. The value of this variable depends on the exponential moving average of current and past squared gradients (up to training step *t*). The meta-parameter *γ* controls the degree of exponential smoothing similarly to *α* of the Momentum method above. Setting *γ* = 0 would be similar to vanilla gradient descent where only the gradient of current training step *t* is used to update. In practice a common heuristic for the choice of *γ* in the deep learning community (also suggested by Tieleman and Hinton (2012)) is 0.999 and instead only tuning the global learning rate *λ*.

At this point we cannot provide a theoretically grounded explanation for the regularizing effect we see and report in the results section when using adaptive learning instead of Momentum. Theoretically grounded explanations of the effects of different gradient-decent optimizers are a very recent and ongoing research field in the Deep Learning community. Even for rate-based artificial neural networks it’s currently not possible to provide a theoretical explanation. We thus conducted an empirical analysis of the weight updates and report our findings in Figure 2C and conclusions in the discussion.

### Detection of spatio-temporal input spike patterns

In this task we study the general case of counting arbitrary, task dependent patterns. To this end we use 1sec long spike trains generated from point processes as a model of complex spatio-temporal patterns that represent features of task dependent activity. An input to the MST model consists of a sequence of such patterns, each of which assigned with a specific target 𝓡*_i_*. The patterns are superimposed onto a 10sec long spike train of background activity. Similar to the task in (Gütig, 2016) the MST model is trained to respond with spikes for each pattern occurrence where the number of spikes per pattern depends on its assigned target 𝓡*_i_*. For each data-set a training set of 200 samples and a separate validation set of 50 samples is generated. Each pattern is associated with a fixed, positive integer target 𝓡*_i_* ∈ [0, 9]. For each data-set the patterns are generated from a different renewal process. Out of the 9 patterns, 5 patterns are considered to be *task-related* and are associated with some positive target 𝓡*_i_*. The remaining 4 patterns are considered to be distractor patterns with target 0. The training target for each of such input spike train is determined as the sum over all individual targets Σ*_i_*𝓡*_i_* of each occurring pattern.

It has been shown that in-vivo cortical spiking activity is typically more regular then Poisson (Mochizuki et al., 2016; Nawrot, 2010). In general any correlated stimuli input is expected to deviate from Poisson (Farkhooi et al., 2011). Moreover, input is generally non-homogenous, i.e. time-varying. However in (Gütig, 2016) only homogeneous Poisson statistic of input patterns and background were considered.

All patterns are generated as 1sec long spike trains by drawing instantaneous firing rates from three different point processes (renewal processes): Γ_1_ representing the homogeneous Poisson process, Γ_5_, and Γ_15_ represent Gamma-Processes (simulated by thinning a Poisson process with 5x and 15x of the Gamma process’ intensity) with a fixed intensity (or rate) of *λ* = 0.89 spike events per second.

Input spike trains of 10 sec duration and 500 presynaptic inputs are generated by simulating 10 sec long spike train of background activity using renewal processes and patterns are superimposed onto this background activity. The number of patterns to appear within a sequence is drawn from a Poisson distribution of mean 5 patterns (with replacement). These patterns are randomly positioned in time within those 10 sec but are not allowed to overlap (an example of an input spike-train is shown in fig. 1A).

We evaluate learning under different noise conditions, where one condition uses homogenous Poisson background activity and the other condition uses inhomogenous Poisson background activity. The homogenous background activity is drawn from a stationary Poisson process (*λ* = 0.3 spikes/sec) while for the inhomogenous case the instantaneous firing rates are slowly modulated by 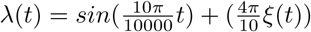 where *ξ*(*t*) is noise drawn from a standard normal distribution.

The free meta-parameters for Momentum and Adaptive learning are set to be *α* = 0.999 and *γ* = 0.999 respectively. These are heuristic values taken from current deep learning frameworks and in practice are treated as constant parameters. Thus, the only real free parameter is the global learning rate *λ*. Since the objective of this task is to study the effect of the two different update methods, we are not concerned to determine the optimal learning rate that would give the best possible, absolute numbers in terms of training error. The described effect in the results section is independent of the specific choice of the learning rate although the absolute numbers vary.

### Counting handwritten digits

This task considers the problem of estimating numerosity. Specifically the problem of counting the occurrences of digit 1 within an image showing 9 random MNIST (LeCun and Cortes, 2010) digits positioned within a 3*x*3 grid. Following Seguí et al. (2015) and Fomoro we generated new images of size 50×50 pixels. Each image is subdivided into a 3*x*3 grid where each grid cell shows a randomly chosen (with replacement), single MNIST digit. Out of the 9 possible cells, up to 6 cells can be occupied by digit 1. This yields samples with possible targets from 0 − 6. The generated data set is only roughly balanced, containing ∼ 200 samples for each target 0 − 6. This imbalances further renders this a difficult task. The model is supposed to learn to count the number of occurrences of the digit by generating 1 output spike each. The training target is provided (supervised learning) by a single scalar label. All models are trained using 5-fold cross-validation. While the training set for the MST model comprises 400 samples, the ConvNet is provided with 800 samples. Additionally, the ConvNet is provided with a much larger learning rate of *λ* = 0.01 to accelerate training, while the MST is manually tuned to use learning rate of *λ* = 0.00002. To train the ConvNet we use the ADAM (Kingma and Ba, 2014) optimizer which has been found to be an effective optimizer for training ConvNets. For the MST model we use our adaptive learning rate method where the meta-parameter is set to *γ* = 0.999. The MST model is trained for max. 30 epochs as it does not improve further after this. The ConvNet is trained for max. 200 epochs. For all models, the training is considered to be converged at that epoch before the validation error diverges for the first time (early-stopping). While the ConvNet shows monotonic decrease of validation error, the MST fluctuates.

For the Multi-Spike Tempotron the images have to be encoded as spike trains. This is done by using *Filter-Overlap Correction Algorithm* (FoCal) (Bhattacharya and Furber, 2010), a 4-layer model of the early visual system that uses an improved rank-order coding originally proposed by Thorpe and Gautrais (1998). Encoding a single 50*x*50px image thus yields a spike train with 4 × 50^2^ = 10000 synapses. The encoding algorithm makes use of spatial correlations in order to reduce the amount of redundant information. This is similar to the convolutional filters embedded in current deep neural networks (Simonyan and Zisserman, 2014; Krizhevsky et al., 2012). For reference, we train a conventional ConvNet architecture that has been shown to successfully accomplish this task when trained on 100000 samples Fomoro. The architecture uses several layers (conv1 - MaxPool - conv2 - conv3 - conv4 - fc - softmax) and includes recently discovered advances like strided and dilated convolutions (Yu and Koltun, 2015).

The free meta-parameters for the MST model, Adaptive learning parameter *γ* and global learning rate *λ*, are set to be *γ* = 0.999 and *λ* = 0.0001. This learning rate has been determined manually, by step-wise decreasing from 0.1 by factor of 10 until reaching the best trade-off between learning speed and convergence of validation error. For the choice of *γ* we refer to the explanation given in the method section above.

### Insect-inspired numerical cognition during visual inspection flights

Following Vasas and Chittka (2019); Howard et al. (2018) we consider estimation of numerosity of geometric shapes during a sequential inspection strategy employed by insects. We use 97 sample trajectories from sequential inspection flights from real honeybees, taken from supplements of Vasas and Chittka (2019). The available trajectories cover samples from 0 to 6 items (we removed 0 since it only had a single trajectory). Following Vasas and Chittka (2019) the trajectories have been used to extract a sequence of single images with a field of view (FOV) of 60° and 2cm distance to the inspected image. Thus each time point of a scanning trajectory yields a 183*x*183 pixel image. Particularly, the absolute difference of each image *S* between two successive time points *t* and *t* + 1 (1st derivative) of the trajectory is used: *FOV_diff_* = |*S*(*t*) − *S*(*t* + 1)|. While the proper way would be to use |*S*(*t* − 1) − *S*(*t*)| we decided to exactly follow the method used in Vasas and Chittka (2019). Differently from Vasas and Chittka (2019) the sequence of derivative images is down sampled to obtain sequences of equal length of 10 images. This is done to reduce the computational cost as well as removing some redundant information from overlapping field of views between two successive time steps (a very coarse approximation of a working memory). All images are further down-scaled by factor 0.25 to 46*x*46 pixels. This additional preprocessing is done solely to reduce computational cost and to reduce the number of free parameters (synapses) in the MST model. To obtain spike trains from the image sequences, each *FOV_diff_* image is encoded as a short parallel spike train using Filter-overlap Correction (FoCal) algorithm (Bhattacharya and Furber, 2010). FoCal resembles a 4-layer early visual system and is an improved rank-order coding scheme of images originally proposed by Thorpe and Gautrais (1998). The resulting parallel spike trains per *FOV_diff_* image are finally concatenated (without gaps) into a single long parallel spike train. Using this encoding results in parallel spike trains with 8468 synapses that converge to the MST. The MST model is trained (supervised) to fit its numbers of output spikes to the precise item count of geometric shapes. We used the adaptive learning method described above with *γ* = 0.999 (also see explanation in methods section above), *λ* = 0.00002 (manually tuned) and performed a 10-fold, stratified cross-validation and trained for max. 25 epochs. We consider a model’s training to be converged at that epoch before the validation error diverges for the first time (early stopping). This generally was the case after 4-7 epochs. We assess the performance on a ‘greater than’ dual choice task following the original experiments of Howard et al. (2018). To this end, we randomly choose two samples of different numerosity (*y*_1_, *y*_2_) (independently for test set & training set) and feed the corresponding images into a randomly chosen, trained instance of the MST (10-fold cross-validation yields 10 independent models in total). A prediction by the MST 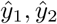 is considered to be correct if 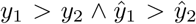 (an vice versa). In undecidable cases where 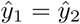 a random decision is made (coin-flip). This sampling process is repeated for 1000 random pairs.

